# Species-informative SNP markers for characterising freshwater prawns of genus *Macrobrachium* in Cameroon

**DOI:** 10.1101/2022.01.24.477480

**Authors:** Judith G. Makombu, Evans K. Cheruiyot, Francesca Stomeo, David N. Thuo, Pius M. Oben, Benedicta O. Oben, Paul Zango, Eric Mialhe, Jules Romain Ngueguim, Fidalis D. N. Mujibi

**Author notes:** currently at The University of Queensland, Australia (Evans K. Cheruiyot). currently at European Molecular Biology Laboratory (EMBL), Heidelberg, Germany. These authors contributed equally (Judith G. Makombu, Evans K. Cheruiyot). **Correspondance**, Dr. Fidalis D. N. Mujibi.

## Abstract

Single Nucleotide Polymorphisms (**SNPs**) are now popular for a myriad of applications in animal and plant species including, ancestry assignment, conservation genetics, breeding, and traceability of animal products. The objective of this study was to develop a customized cost-effective SNP panel for genetic characterisation of *Macrobrachium* species in Cameroon. The SNPs identified in a previous characterization study were screened as viable candidates for the reduced panel. Starting from a full set of 1,814 SNPs, a total of 72 core SNPs were chosen using conventional approaches: a) allele frequency differentials, minor allele frequency (**MAF**) profiles, and Wright’s Fst statistics. The discriminatory power of reduced set of informative SNPs were then tested using the admixture analysis, principal component analysis (**PCA**), and discriminant analysis of principal components (**DAPC**). The panel of prioritised SNP markers (i.e., N = 72 SNPs) distinguished *Macrobrachium* species with 100% accuracy. However, large sample size is needed to identify more informative SNPs for discriminating genetically closely related species, including *M. macrobrachion* versus *M. vollenhovenii* and *M. sollaudii* versus *M. dux*. Overall, the findings in this study show that we can accurately characterise *Macrobrachium* using a small set of core SNPs which could be useful for commercial breeding operations, conservation, and species assignment of this economically important species in Cameroon. Given the results obtained in this study, a larger independent validation sample set will be needed to confirm the discriminative capacity of this SNP panel for wider commercial and research applications.

## Introduction

Freshwater prawns of the genus *Macrobrachium* Bate, 1868 (Crustacea, Decapoda, Palaemonidae) are a highly diverse group of decapod crustaceans of high economic importance globally. They occur in diverse habitats worldwide, from brackish estuarine to upland streams of the tropics and subtropics (Holthuis, 1980;March et al., 2002). A total of 240 *Macrobrachium* species are presently known (Holthuis, 1980;Wowor et al., 2009;De Grave and Fransen, 2011), although it is difficult to estimate the correct species richness of *Macrobrachium*, as new taxa are often described every year.

Ecologically, *Macrobrachium* plays a critical role in stream food webs because it serves as intermediate consumer, linking the production of periphyton and detritus with higher trophic groups (Browder et al., 1994). Economically, *Macrobrachium* serves as important food resources for carnivorous fish and humans, it is amongst the main target species for fisheries and aquaculture (Bowles et al., 2000) and it sustains most viable artisanal and commercial fisheries in West Africa Sub-region (Okogwu et al., 2010). However, the *Macrobrachium* fauna of West Africa is poorly understood.

The main taxonomic records date back to the general investigations of decapod crustaceans in West Africa (Monod, 1980;Powell, 1980), which reported 10 species of *Macrobrachium* in this region including four in Cameroon. Recent studies pointed out the higher species richness of Cameroonian *Macrobrachium* (Doume et al., 2013;Siméon et al., 2014;Makombu et al., 2015) and increased the number of known species from four to six: *M. vollenhovenii*, *M. macrobrachion*, *M. chevalieri*, *M. sollaudii*, *M. dux*, *M. felicinum*. All these studies used morphological keys. It is well-known that morphological identification of species of this genus is quite difficult because many features used for identification are common to all known species (Zhang et al., 2009). These studies illustrate that traditional morphological characters alone are insufficient in the accurate diagnosis of the genus *Macrobrachium*. Molecular data has proven very useful to elucidate the taxonomic relationships in morphologically variable groups of freshwater prawns (Vergamini et al., 2011). Several studies have used mitochondrial DNA sequence data from the 16S rRNA and cytochrome c oxidase subunit 1 (CO1) genes to characterize Asian *Macrobrachium* taxonomy, biogeography, evolution, and life history (e.g., Murphy and Austin, 2003;Zhang et al., 2009;Vergamini et al., 2011). Microsatellite markers have also been developed for *Macrobrachium rosenbergii* (Divu et al., 2008).

In our recent work (Makombu et al., 2019), we used Diversity Arrays Technology (DArT) (Kilian et al., 2012) to genotype and characterize *Macrobrachium* species from the coastal area of Cameroon using 1,814 SNPs. In that study, we identified at least four species of *Macrobrachium* based on the ADMIXTURE analysis and five species when using principal component analysis (PCA), using 1814 SNP markers and for 93 individuals from different species initially differentiated using morphological keys. In this study, we set out to identify a smaller set of informative SNP markers that can be used to characterize *Macrobrachium* species, with the aim of ultimately reducing the cost of genotyping to allow a larger number of individuals to be evaluated in future studies. This is in line with similar studies in humans (Pakstis et al., 2007), wildlife (Oliveira et al., 2015), livestock (Bertolini et al., 2015;Schiavo et al., 2020), and crops (Nguyen et al., 2020). Pakstis et al., (2007) screened 432 SNPs and chose 40 informative SNP markers for forensics and paternity testing in humans. Similarly, Oliveira et al. (2015) screened a total of 158 SNPs and identified a suite of 35 SNPs for genetic inference of domestic cats and European wildcats, while Bertolini et al. (2015) identified 48 and 96 SNPs from a set of 50K SNPs for breed assignment in cattle. Besides the cost, the informative set of markers for *Macrobrachium* species could be useful for routine use in species ancestry assignment, conservation, forensics, and breeding purposes.

Several methods have been proposed for identification of informative genetic markers for inference of population structure, such as the Delta method, which estimates allele frequency difference between pairs of populations (Shriver et al., 1997), Fst variants [e.g., Weir & Cockerham (Weir and Cockerham, 1984)], informativeness for assignment (*I*_n_) (Rosenberg et al., 2003), and PCA (Price et al., 2006). These methods are closely related and gives comparable results (Pfaffelhuber et al., 2020). More recently, Schiavo et al. (2020) used a machine learning approach (Random Forest) to select 96 informative SNPs from the 60K porcine array for use in discriminating pig breeds.

In this study, we chose a smaller set of informative SNPs for genetic characterisation of *Macrobrachium* species from a full set of 1,814 SNPs using conventional approaches: a) SNPs with high Weir & Cockerham Fst values (Weir and Cockerham, 1984), and b) SNPs defined as ‘private’ or unique for each study species because they are segregating (i.e., not fixed) in only one out of the seven populations studied.

## Materials and Methods

### Samples, genotyping and quality checks

The dataset used in this study is part of our previous work and has been described in more detail by Makombu et al. (2019), including the sampling locations, morphological characterization, and genotyping. Briefly, we collected a total of 1,566 *Macrobrachium* specimens from fishermen catches between May 2015 and April 2016 from different regions in Cameroon: Lokoundje, Kienke, and Lobe Rivers, in the South region; at Batoke, Mabeta and Yoke rivers in the South West region and Nkam and Wouri rivers in the Littoral region of Cameroon [comprehensive descriptions of the specimens were given by Makombu et al. (2019)]. Out of these samples, a small set of 93 individuals was selected for genotyping representing seven species: 18 samples from *M. dux*; 18 *M. macrobrachion*; 18 *M. sollaudii*; 17 *M. vollenhovenii*; 12 *M. chevalieri*; 5 *M. felicinum*, and 5 *M*. sp (an undescribed species). These species were identified based on the morphological key described by Monod (1980) Monod (1980) and Konan et al. (2008). The images of the study species are shown in Figure 1.

**Figure 1.**
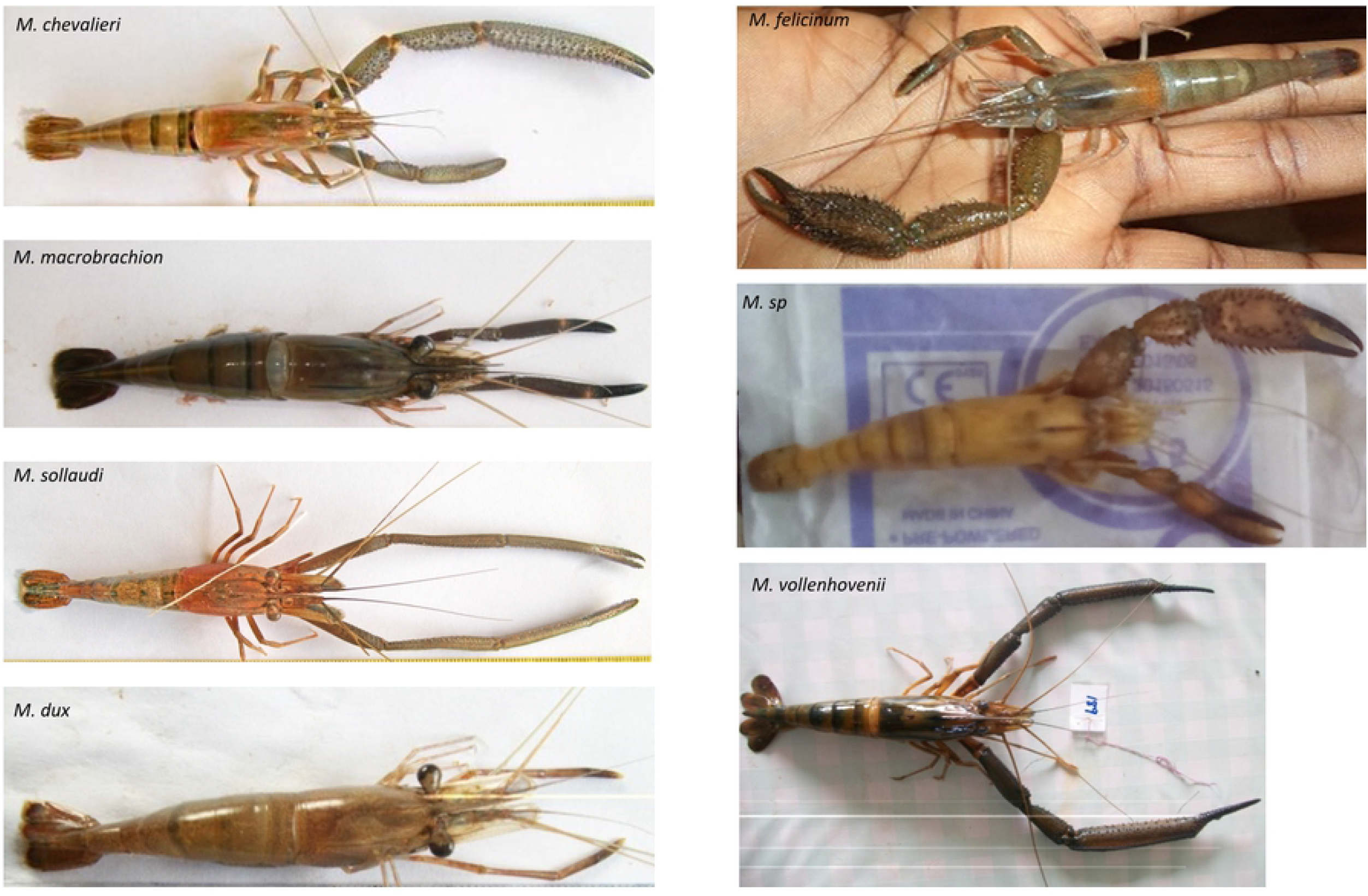
Pictures of seven *Macrobrachium* species in the coastal area of Cameroon identified based on morphological analysis using Monod (1980) and Konan (2008) key.

Following DNA extraction and genotyping using DArT markers (Kilian et al., 2012), a total of 1,814 out of 52,834 SNPs were retained for data analysis, as described in Makombu et al., (2019). This remaining set of markers (N = 1,814) passed quality checks based on call rate > 80% and minor allele frequency (MAF > 5%). We used this set of markers in this study as a benchmark to test species assignment to the respective population versus the reduced set of ‘informative SNPs’.

### Identifying informative SNP markers

We used the following steps to identify ‘private’ SNPs (i.e., those segregating in only one species) for *Macrobrachium* species [*M. dux*, *M. macrobrachion*, *M. sollaudii*, *M. vollenhovenii*, *M. chevalieri; M. felicinum*, *M*. sp]:

1. Compute allele frequencies for each population and SNP (N = 1,814) using the Hierfstat package (Goudet, 2005) in R (R Core Team, 2018).
2. Select SNPs that are segregating in only one population (i.e., fixed allele frequencies in six out of seven species or populations studied).
3. For the SNPs in set 2 above, select SNPs segregating with a minimum threshold of 0.03 for the alternate allele to avoid fixed SNPs.
4. Repeat the above steps (i.e., 1 to 3) for 100 runs by randomly selecting 80% of the individuals in each species for each repeat run.
5. For private SNPs in step 4, select two sets of SNPs: a) informative or ‘private SNPs’ identified in > 50% (i.e., > 50 runs) of the 100 repeated runs, and b) informative or ‘private SNPs’ identified in > 80 runs (considered as most stable core SNPs). These SNP sets will be called ‘private SNPs 50’ and ‘private SNPs 80’ panels.

As an alternative approach, we computed Weir & Cockerham Fst values (Weir and Cockerham, 1984) for a full set of SNPs (N = 1,814) using PLINK software v1.9 (Purcell et al., 2007). We then selected SNPs with relatively high Fst values (> 0.7; Supplementary Figure 1). Most of the SNPs with high Fst values overlapped with those identified in the first approach [i.e., step 5(b) above], except for a few SNPs (N = 9) with low Fst values (meaning less informative SNPs), all of which were from *M. chevalieri*. Therefore, we excluded these SNPs (N = 9) and focused analysis on the ‘private SNPs’ or those considered as the most informative SNPs identified using the first approach (i.e., private SNPs). Besides, this species (i.e., *M. chevalieri*) is the most genetically divergent (see Figure 4), suggesting that a relatively few core SNPs are needed to distinguish from other *Macrobrachium* species. For the selected set of informative SNPs, we calculated allele frequency (MAF), observed, and expected heterozygosity for each population using the Hierfstat package (Goudet, 2005) in R (R Core Team, 2018). An overview of the SNP identification and validation is described in Figure 2.

**Figure 2.**
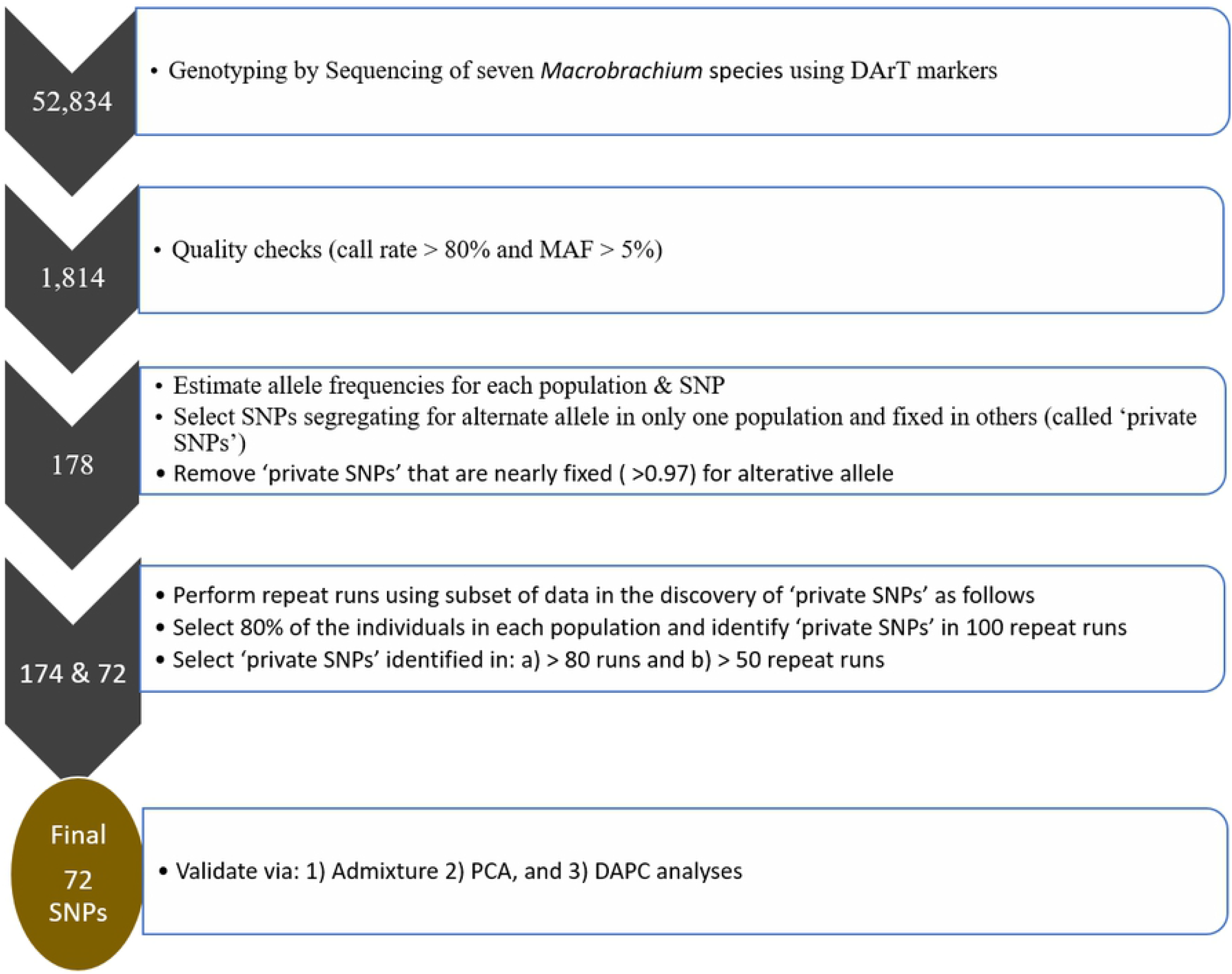
An overview of the identification and validation of ‘private SNP’ or informative SNPs. PCA – principal component analysis; DAPC – discriminant analysis of principal components; MAF – minor allele frequency.

### Validation of informative SNPs

We used three approaches to test whether selected ‘private SNPs’ are parsimonious or robust in discriminating *Macrobrachium* populations: a) principal component analysis (PCA) using PLINK software v1.9 (Purcell et al., 2007), b) discriminant analysis of principal component (DAPC), and admixture analysis both (i.e., b and c) using the Adegenet package (Jombart, 2008) in R (R Core Team, 2018). Notably, PCA and DAPC methods are comparable except that the former aims to discern the overall variability in the population (i.e., within- and between-group variability), while the latter focuses on distinguishing between-group components (Jombart et al., 2010). Using the whole SNP set as a benchmark, we tested population assignment with two sets of informative SNPs:

a. ‘private SNPs’ identified in step 5 (a) above (i.e., those identified in > 50 times of the repeated random subsets – ‘private SNPs 50’).
b. ‘private SNPs’ from step 5 (b) above (i.e., those detected in >80 of the repeated random subsets – ‘private SNPs 80’). However, most of the ‘private SNPs Full’ panel overlapped with the ‘private SNPs 50’ panel (i.e., 178 out of 174 SNPs). Therefore, we only tested ‘private SNPs 50’ (N = 174 SNPs) and ‘private SNPs 80’ (N = 72 SNPs) against a benchmark marker set (N = 1,814 SNPs).

## Results and Discussion

While the cost of genotyping has reduced considerably over the years, thanks to the rapid evolution of high-throughput technologies, it was not feasible to cost-effectively genotype a large population of highly diverse species such as *Macrobrachium* using dense genetic markers in our previous work (e.g., Makombu et al., 2019). The objective of this study was to identify and test the effectiveness of a small set of SNPs for characterising *Macrobrachium* species. Consequently, we have demonstrated that it is possible to accurately discriminate between *Macrobrachium* species using a smaller suite of highly informative SNPs (N = 72). These findings are promising, meaning if validated in an independent population, we can genetically screen a large population of *Macrobrachium* at a lower cost using informative variants. This can facilitate forensics, conservation, and breeding purposes for this economically important species, particularly for resource-poor farmers in developing countries.

We used several conventional statistics to choose a small set of highly informative SNPs for characterising *Macrobrachium* species: a) private SNPs (Phillips et al., 2007) – defined as those segregating in only one population and fixed in others b) minor allele frequency, and c) SNPs with high Fst values > 0.70. We then validated prioritised SNPs using empirical (i.e., admixture analysis) and heuristic (PCA and DAPC) approaches. Overall, we found that the reduced set of 72 informative SNPs can classify *Macrobrachium* individuals into respective populations with 100% probability based on the ADMIXTURE results. Similarly, the PCA and DAPC methods showed good agreement when comparing clustering profiles of *Macrobrachium* species obtained from using a full set of SNPs (N = 1,814) versus a reduced set of informative SNPs (N = 72).

### Minor allele frequency (MAF)

MAF is an important metric for evaluating the informativeness of genetic variants and has been used to develop custom SNPs arrays in cattle and other species (e.g., Matukumalli et al. (2009)). Figure 3 shows the distribution of MAF for *Macrobrachium* populations that were computed separately for each population. Most of the SNPs have low minor allele frequency (i.e., MAF < 0.1) across populations. However, a sizable number (N = 243) have relatively high MAF values (MAF > 0.1). Notably, the high proportion of SNPs with low MAF (i.e., < 0.1) was expected since the *Macrobrachium* genome is still poorly annotated. This is comparable to other studies in cattle (e.g., Cheruiyot et al. (2018)) that reported a larger proportion of SNPs with low MAF for indicine breeds (less genetically described breed) compared to well-known Holstein breeds.

**Figure 3.**
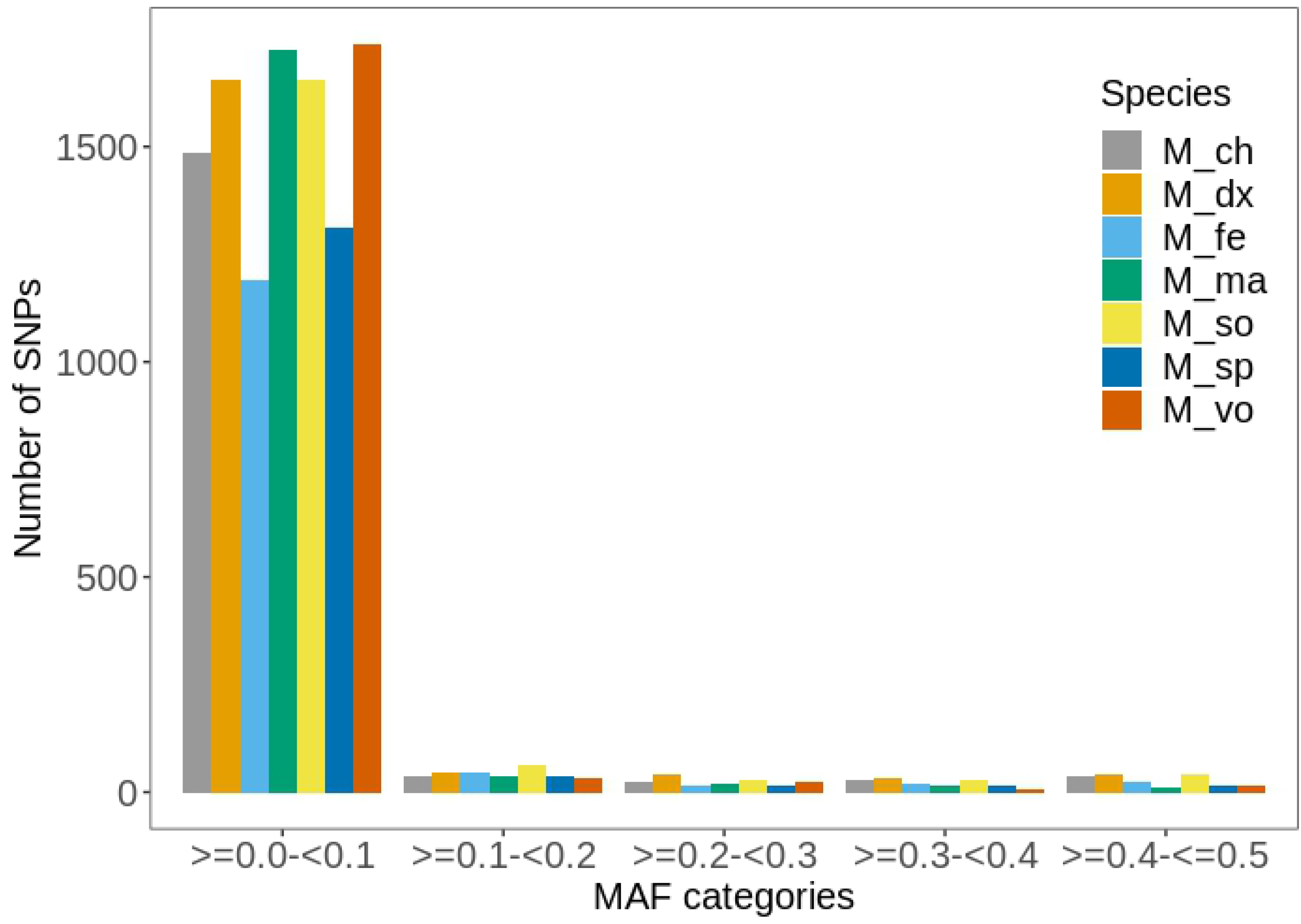
Distribution of minor allele frequency (MAF) for different *Macrobrachium* species: *M. dux*, *M. macrobrachion*, *M. sollaudii*, *M. vollenhovenii*, *M. chevalieri; M. felicinum*, and *M. sp*.

### Species-informative ‘private SNPs’

Table 1 shows the summary statistics for the final set of ‘private SNPs’ (N = 72) identified in this study from a starting full set of 1,814 SNPs. Notably, these SNPs (i.e., N = 72) represent those identified from repeated re-sampling analysis (‘private SNPs 80’; see Methods) considered stable or of high-quality; therefore, more relevant for species population assignment. The number of ‘private SNPs’ ranged from 2 (*M. dux*) to 16 (*M. chevalieri*). The fact that we found only 2 ‘private SNPs’ for *M. dux* is not surprising considering that this species appears to be genetically closely related to *M. sollaudii* species based on phylogenetic analysis (Figure 4). The same case applies to *M. vollenhovenii* and *M. macrobrachion* – also genetically closely related species (Figure 4). While a possible reason for this close genetic relationship could be because of gene flow, our admixture results (Figures 7 and 8) suggest very limited admixture among these species. Alternatively, a more plausible reason could be that these species [i.e., *M. vollenhovenii* versus *M. macrobrachion* and *M. sollaudii* versus *M. dux*] are conspecific, meaning that classifying them as separate species using morphological keys could be misleading. We found most of the *M. sollaudii* samples were males, whereas *M. dux* were mainly females [see Makombu et al. (2019)]. Notably, the few *M. sollaudii* individuals classified as females were all young or juveniles. Overall, these observations suggest that it is highly likely that the morphological key is perhaps separating males and females of the same species.

**Table 1.** Summary statistics of the ‘private SNPs’ (‘private SNPs80’; N = 72) identified in the study.

| Population | N samples | 'private |  | Obs | Exp | Fst |
| --- | --- | --- | --- | --- | --- | --- |
|  |  | SNPs'* | MAF<sup>□</sup> | (Het) | (Het) | values‡ |
| *M. chevalieri* | 13 | 16 (60) | 0.05 - 0.48 (0.12) | 0.073 | 0.076 | 0.75 – 0.91 |
| *M. dux* | 18 | 2 (23) | 0.38 - 0.38 (0.38) | 0.002 | 0.002 | 0.98, 0.96 |
| *M. felicinum* | 5 | 14 (24) | 0.05 - 0.41 (0.19) | 0.09 | 0.079 | 0.83 – 0.98 |
| *M. macrobrachion* | 17 | 14 (26) | 0.05 - 0.47 (0.26) | 0.01 | 0.016 | 0.91 – 0.98 |
| *M. sollaudii* | 18 | 8 (17) | 0.05 - 0.38 (0.31) | 0.008 | 0.009 | 0.85 – 0.98 |
| *M. sp* | 5 | 11 (16) | 0.07 - 0.43 (0.26) | 0.062 | 0.057 | 0.84 – 0.98 |
| *M. vollenhovenii* | 17 | 8 (7) | 0.06 - 0.47 (0.27) | 0.007 | 0.006 | 0.85 – 0.98 |

\*The 'private SNPs' in brackets represents those identified from the less stringent repeated cut-off after repeat runs (i.e., 'private SNPs50'; N = 174); ‡Fst values were computed based on Weir & Cockerham (1984) using PLINK software (Purcell et al., 2007); MAF – minor allele frequency; <sup>□</sup>Values in brackets represents average MAF.

**Figure 4.**
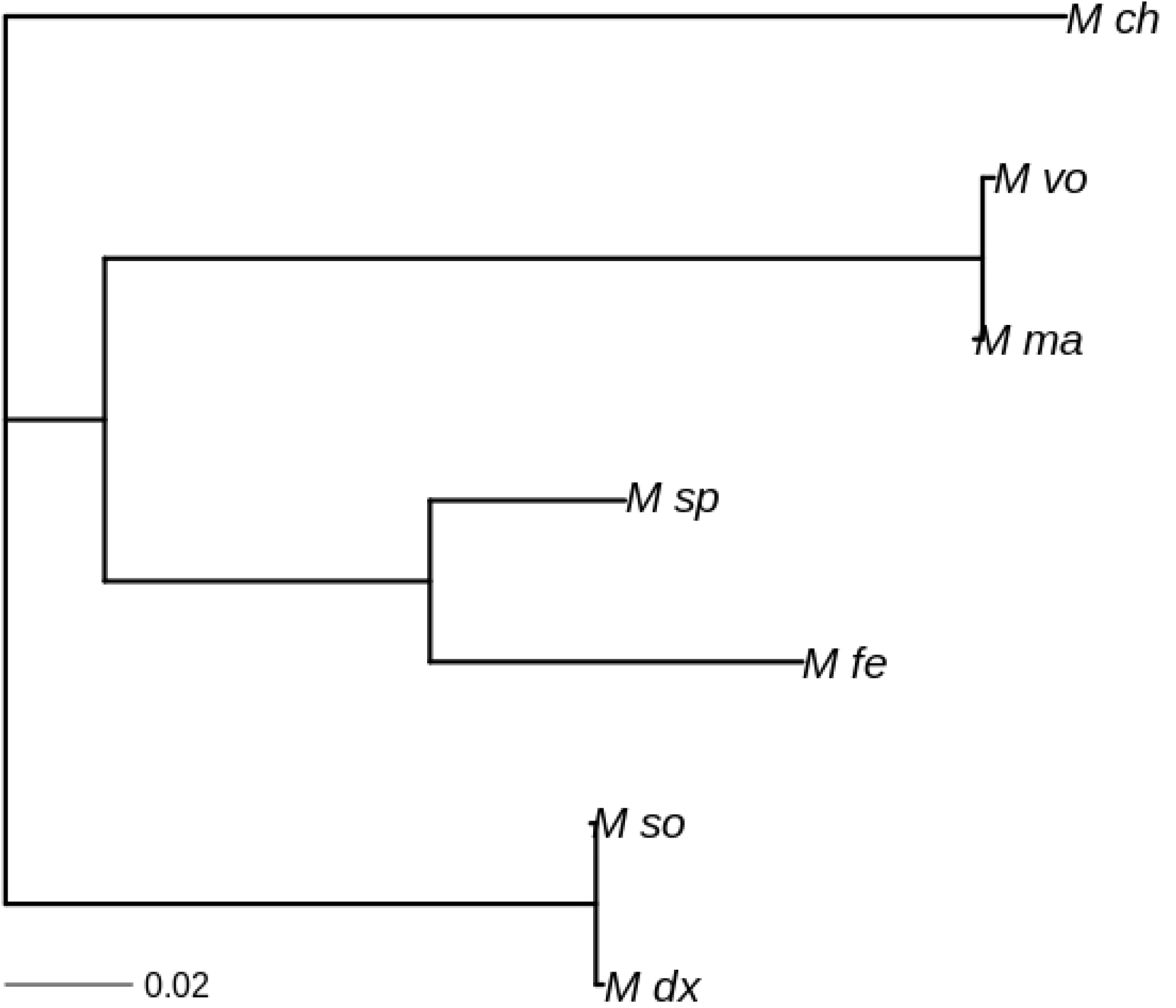
Phylogenetic tree obtained using 72 private SNPs for *Macrobrachium* species: *M. dux*, *M. macrobrachion*, *M. sollaudii*, *M. vollenhovenii*, *M. chevalieri; M. felicinum*, *and M. sp*.

Therefore, future studies with large sample sizes are needed to conclusively determine if these closely related individuals belong to the same species.

If indeed these species (i.e., *M. sollaudii* versus *M. dux*) are separate but with similar genetic relationship, then it means that many SNPs are required to discriminate between species. In contrast, we need a smaller number of informative SNPs to distinguish *M. chevalieri* versus other species, given that this species is genetically divergent compared to other species (Figure 4). This is consistent with the work of Wilkinson et al. (2011) in which the authors show that relatively more SNPs are required to characterise closely related cattle breeds. As such, we recommend further work with a larger sample size to identify more core SNPs, particularly for closely related *Macrobrachium* species identified in this study.

To date, the domestication and commercial aquaculture of *Macrobrachium* prawns have not been successful in Africa, unlike other species such as *M. rosenbergii*, which is widely cultured in other parts of the world (New et al., 2009). However, work is currently underway in Cameroon to breed *M. vollenhovenii* as a food resource for humans (J. Makombu; personal communication) and as a biocontrol species for Schistosomiasis – a serious parasitic disease affecting humans (Savaya-Alkalay et al., 2018). *M. vollenhovenii* species is often preferred for aquaculture because the adults are usually bigger compared to other *Macrobrachium* prawns. An attempt to crossbred *M. vollenhovenii* and *M. rosenbergii* by Savaya-Alkalay et al. (2018) was unsuccessful. Interestingly, in our field sampling, we found some *M. macrobrachion* adult individuals of the same size as *M. vollenhovenii* species. As noted earlier, we think that these two species are conspecific. This is supported by the phylogenetic tree (Figure 4) and the admixture results (Figure 7). While we identified 14 and 8 informative SNPs for *M. macrobrachion* and *M. vollenhovenii* species, respectively, it may be necessary to consider a smaller one set of core SNPs for characterising these species, if it is conclusively established that they are indeed the same species of *Macrobrachium*.

The average MAF calculated from the ‘private SNPs’ (N = 72) [based on a combined dataset for all *Macrobrachium* species] ranged from 0.12 (*M. chevalieri*) to 0.38 (*M. dux*). Similarly, the observed and expected heterozygosity values were low, with the average estimates of 0.036 and 0.035. On the other hand, the Fst values were high (> 0.7) for this ‘private SNP’ set. The high Fst (> 0.7) and MAF (i.e., > 0.1) cut-off for ‘private SNPs’ chosen in this study suggests that they are highly informative for characterising *Macrobrachium* species.

Other studies have also identified core marker sets for characterising various species, including humans (Pakstis et al., 2007), cattle (Bertolini et al., 2015), wildlife (Oliveira et al., 2015), and plants (Nguyen et al., 2020). For example, Bertolini et al. (2015) chose 48 and 96 informative SNPs for cattle from the 50k SNP chip based on the principal component analysis and machine learning methods (random forest). In recent work, Schiavo et al. (2020) followed a similar approach as Bertolini et al. (2015) and identified a small set of informative SNPs for pigs from the porcine 60k array. While we discovered a total of 72 ‘private SNPs’ in this study, even a smaller number of high-quality SNPs is desirable to minimize genotyping costs for *Macrobrachium* species. However, a larger sample size is needed for an informative panel to be developed.

## Validation of informative SNPs

### PCA and DAPC using full set of SNPs

We used the results from the full set of SNPs for PCA and DAPC analysis as a benchmark to see how well different species are classified compared to the reduced set of core markers. Figure 5 shows the PCA and DAPC plots obtained when using a full set of SNPs (N = 1,814). The PCA plot in this study mirrors that reported by Makombu et al. (2019), in which five *Macrobrachium* populations were reported with the following clusters: *M. dux* and *M. sollaudii* (cluster 1); *M. macrobrachion* and *M. vollenhovenii* (cluster 2); *M. chevalieri* (cluster 3); *M. felicinum (cluster 4); M. sp* (cluster 5). This compares well with the results from the DAPC analyses when assuming 5 clusters of *Macrobrachium* species (Figure 5). In addition, these results are consistent with those from phylogenetic analysis discussed earlier (Figure 4). This phylogenetic profile is consistent with the one reported by Makombu et al. (2019) using a large set of 1,814 SNPs. Notably, these plots will be used as the basis to compare how well clustering performs when using the reduced set of ‘private SNPs’.

**Figure 5.**
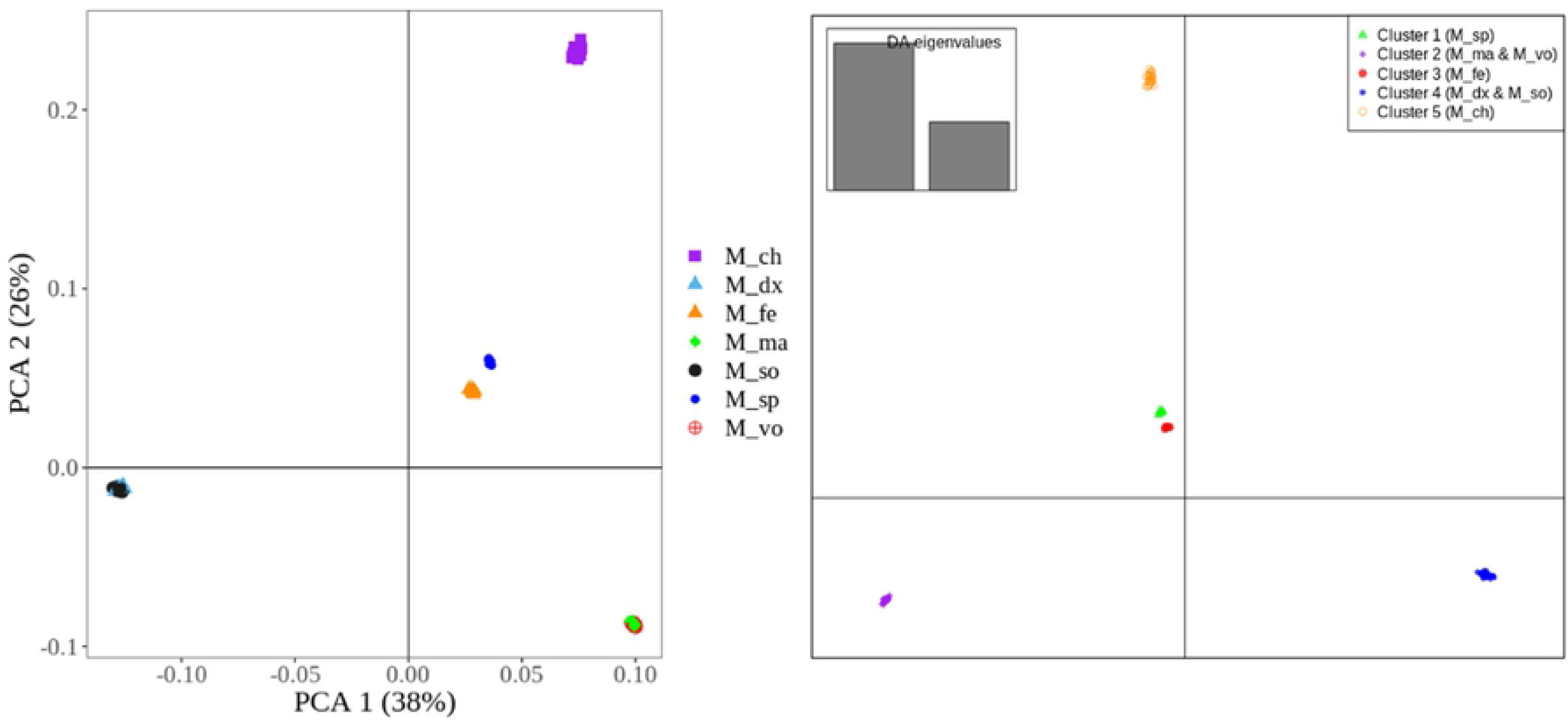
PCA (left plot) and DAPC (right plot) obtained from using a full set of SNPs (N = 1,814). PCA – principal component analysis; DAPC – discriminant analysis of principal components. *M_ch* – *M. chevalieri;M_dx* – *M. dux;M_fe* – *M. felicinum; M_ma* – *M. macrobrachion; M_so* – *M. sollaudii; M_sp* – *M. sp; M_vo – M. vollenhovenii*.

### PCA and DAPC using informative SNPs

Figure 6 shows the PCA and DAPC plot obtained from the reduced set of ‘private SNPs’ (N = 72) considered as more stable or high-quality (i.e., the ‘private SNPs’ called ‘private SNPs 80’; see Methods). For PCA, these SNPs clearly distinguished four groups of *Macrobrachium* species with *M. felicinum* and *M. sp* appearing as one cluster, which contrast with the results obtained from the using full set of SNPs (as described above). However, the plot for PC1 versus PCA3 using ‘private SNPs 80’ clearly separated these species into two distinct populations (Supplementary Figure 1), indicating a total of five *Macrobrachium* species.

**Figure 6.**
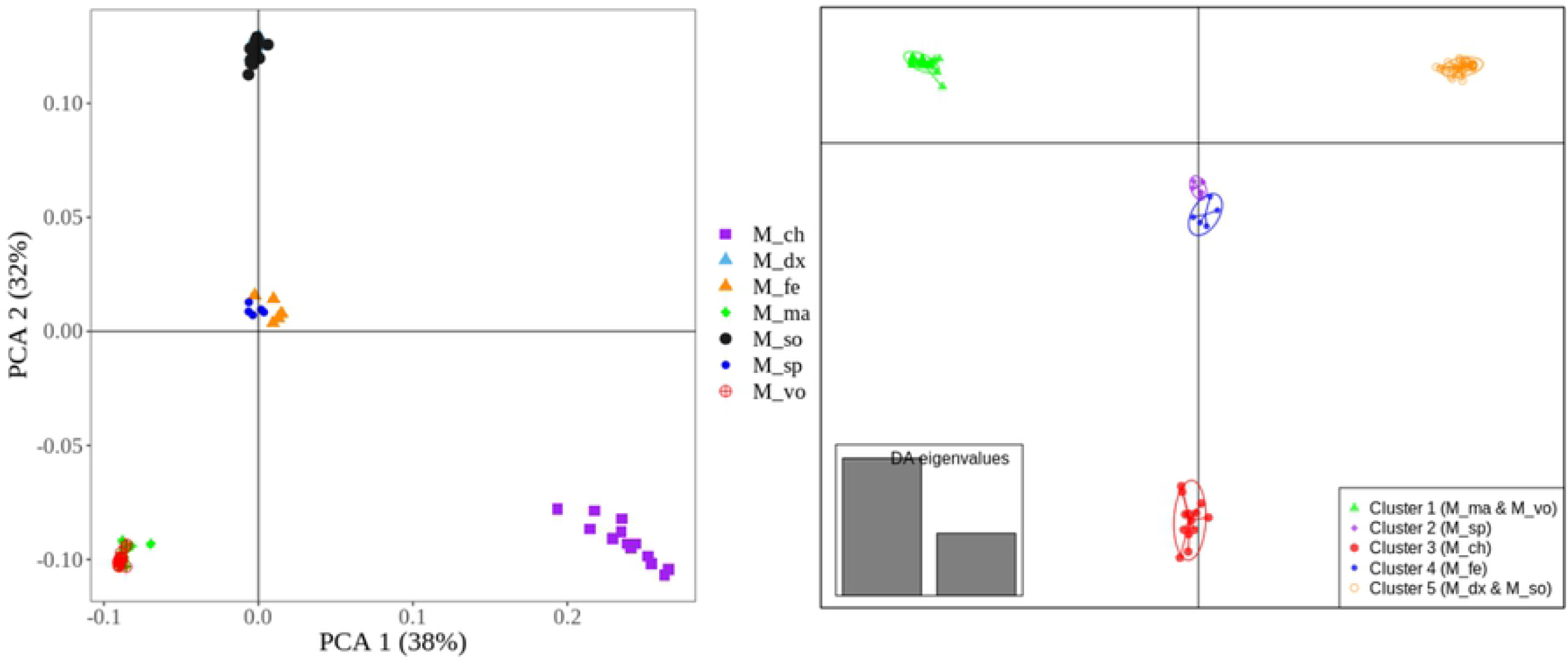
PCA (left plot) and DAPC (right plot) obtained from using the reduced set of ‘private SNPs’ (N = 72 SNPs, called ‘private SNPs80’ panel; see Methods). PCA – principal component analysis; DAPC – discriminant analysis of the principal components. *M_ch* – *M. chevalieri; M_dx* – *M. dux; M_fe* – *M. felicinum; M_ma* – *M. macrobrachion; M_so* – *M. sollaudii; M_sp* – *M. sp; M_vo* – *M. vollenhovenii*.

The above results from PCA (Figure 5), are consistent with those from the DAPC method (Figure 6) where *Macrobrachium* species were clustered into five populations. Notably, the individuals in Figure 5 above (i.e., benchmark results obtained from using the full set of SNPs, N =1,814) appear as tightly clustered within their respective groups compared to those observed when using the ‘private SNPs’ in Figure 6. Overall, these results suggest that a small set of core SNPs can accurately separate *Macrobrachium* populations. Nonetheless, an obvious limitation of our study is the fact that the population used to discover informative SNPs and validation of this SNP set were the same. As such, future work using an independent sample is needed to confirm the discriminatory power of the selected core SNPs.

### Admixture/membership classification using the full set of SNPs

Apart from PCA and DAPC, we also performed admixture analysis to validate the prioritised set of SNPs. Figure 7 shows the admixture results obtained from using the full set of SNPs (N = 1,814), which we considered as the benchmark for subsequent analyses using the reduced set of ‘private SNPs’. The Bayesian information criterion (BIC) plot showed at least 4 to 6 populations of *Macrobrachium* species in the dataset based on the line of deflection in Figure 7. When assuming four groups (K = 4) as the optimal representation of the species in the dataset, we found that all the individuals clustered into their respective groups with 100% probability (Figure 7). This is comparable to the work of Makombu et al. (2019) when assuming the same K value (i.e., K = 4). These results also mimic those obtained from DAPC (assuming four clusters) analysis described earlier (Figure 5). By looking at Figure 5, *M. felicinum* and *M. sp* were separated into different groups when assuming K = 5, which somewhat differs from the results of Makombu et al. (2019), where these two species remained as one group at K = 5, most likely due to the different methods used for admixture analyses. Here, we used the Adegenet program by Jombart (2008), while Makombu et al. (2019) used the ADMIXTURE program (Alexander et al., 2009). The Adegenet program uses discriminant analysis of principal components (DAPC) to infer population clusters, while the ADMIXTURE program applies the Bayesian clustering method. Another difference between the two programs is that the Adegenet uses the K-means algorithm and model selection to find the optimal number of clusters. In contrast, the ADMIXTURE program requires a *priori* definition of the best number of clusters in a dataset. The Adegenet program is designed to maximise between-group difference over within-group difference (Jombart et al., 2010). Regardless of the program used in the analysis, it is important to note, however, that the results from this work and those from Makombu et al. (2019) were generally comparable.

**Figure 7.**
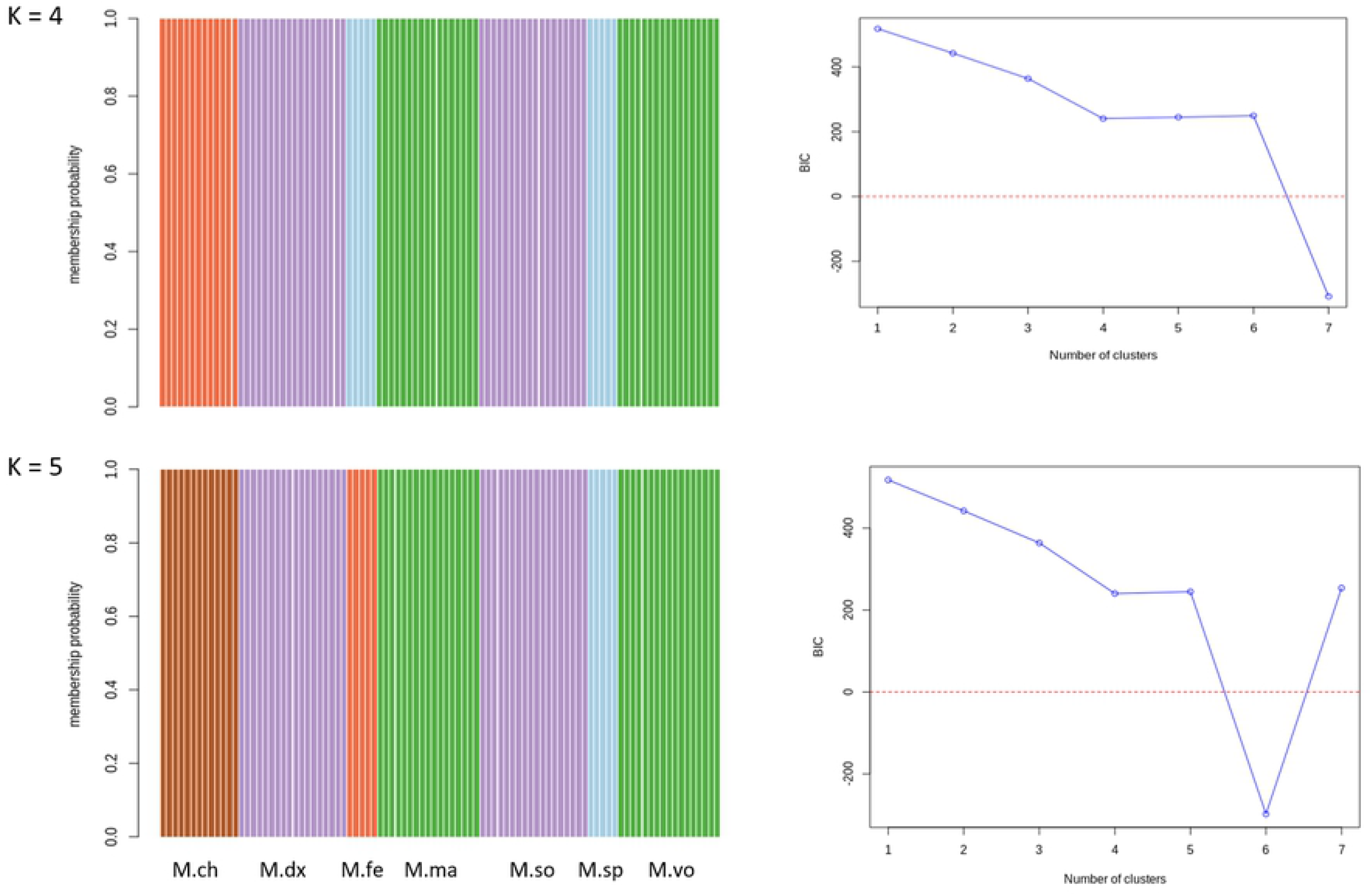
Cluster membership classification of *Macrobrachium* species and the Bayesian information criterion (BIC) plot obtained from the full set SNPs (N = 1,814) using the Adegenet package (Jombart, 2008) assuming four (K = 4) and five (K = 5) populations. Each bar in the admixture plot (left) represents an individual: M.ch – *M. chevalieri;* M.dx – *M. dux;* M.so – *M. sollaudii;* M.fe – *M. felicinum;* M.sp – *M. sp; M.ma* – *M. macrobrachion;* M.vo – *M. vollenhovenii*.

### Admixture/membership classification using reduced set of ‘private SNPs’

Figure 8 shows the admixture results obtained from using a reduced set of ‘private SNPs’ (N = 72 SNPs) described in Table 1. The BIC plot clearly shows that assuming five populations is the most parsimonious to the dataset based on the point-of-line deflection. We, therefore, used K = 5 to display admixture proportions for each sample (Figure 8). By looking at the admixture results, most individuals were classified into their distinct groups (N = 5) with 100% probability, except for a few admixed individuals within *M. felicinum* and *M. sp* populations. These results are consistent with those found when using the full set of SNPs (N = 1,814) described above (Figure 5) and those from the PCA and DAPC analyses (Figure 4). The ADMIXTURE results obtained from the reduced set of 72 core SNPs were also comparable with those from another set of ‘private SNPs’ panel (N = 174; see Methods for description) (Supplementary Figure 2).

**Figure 8.**
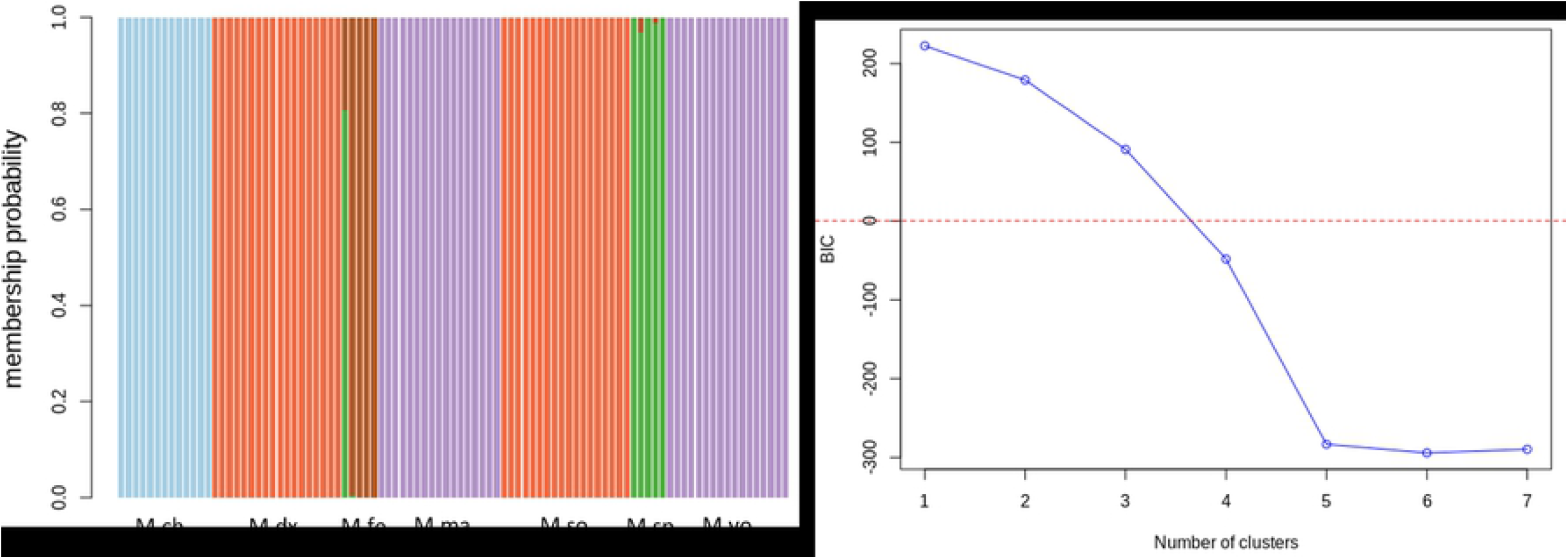
Cluster membership classification of *Macrobrachium* species and the Bayesian information criterion (BIC) plot obtained from a reduced set of informative or ‘private SNPs’ (N = 72 SNPs) using Adegenet package (Jombart, 2008) assuming five clusters (K = 5). Each bar in the admixture plot (left) represents an individual: M.ch – *M. chevalieri;* M.dx – *M. dux;* M.so – *M. sollaudii;* M.fe – *M. felicinum;* M.sp – *M. sp;* M.ma – *M. macrobrachion;* M.vo – *M. vollenhovenii*.

For comparison, we also assessed ancestry classification using the ADMIXTURE software (Alexander et al., 2009) for the reduced set of ‘private SNPs’ (N = 72). Consequently, we found consistent results with those from the Adegenet package (Jombart, 2008) [which is based on the discriminant analysis], when assuming K = 5 with the ADMIXTURE software. As seen in the Supplementary Figure 3 (i.e., results from the ADMIXTURE software), the *Macrobrachium* species were clustered into five groups (at K = 5) with almost 100% ancestry probability in each group.

As discussed earlier, we have used conventional methods (MAF, Fst, and allele frequency differentials) to select informative SNPs for *Macrobrachium* species. Alternatively, more advanced methods such as machine learning (e.g., Bertolini et al., 2015) could be used to identify core SNPs for *Macrobrachium* species. However, the fact that the genome for *Macrobrachium* is still poorly annotated makes it difficult to apply such methods. For example, in this study most individuals had missing genotypes in one or more SNPs. Similarly, more recent supervised methods (e.g., Pfaffelhuber et al., 2020) rely on linkage disequilibrium (LD) which is not possible to apply for our study species with poorly annotated genomic map. Notably, our SNP dataset lacked chromosomal positions. Nevertheless, efforts are underway in several countries to map the complete genome of *Macrobrachium* species, e.g., Jin et al. (2021).

We used group re-sampling approach to select high-quality or stable set of SNPs for characterising *Macrobrachium* species and to guard against false positives (see Methods). However, the small sample sizes within each *Macrobrachium* species makes such re-sampling efforts less effective, meaning a large sample set is needed to identify new informative SNPs and confirm our results. In addition, methods which combine group resampling and machine learning approaches (e.g., Pardy et al., 2011) could be tested in future studies for *Macrobrachium* species. A new validation dataset will have to be provided to test these approaches and the discriminating power of the selected SNP panels.

The core SNPs chosen in this study could be extremely useful in the genetic characterisation of cryptic species of *Macrobrachium*. In our previous work (Makombu et al., 2019), we identified a potentially new species of *Macrobrachium*, which we named *M. sp* (Figure 4). This species is morphologically closely related to another species of *Macrobrachium* called *M. felicinum* (see the images in Figure 1). Considering that *M. sp* was identified from only a few genotyped samples (N < 20) with large sample sizes availed, it is highly likely that new species are yet to be discovered. As such, the core SNPs from this study could facilitate the cost-effective screening of thousands of *Macrobrachium* individuals to identify new species for breeding purposes. In addition, given the current scenario of climate changes, the findings of this study can facilitate documenting new *Macrobrachium* species that are potentially at risk of extinction to inform conservation efforts before they are lost.

## Conclusion

Overall, the results in this study show that we can use a small set of 72 highly informative SNPs to characterise *Macrobrachium* species from the coastal area of Cameroon with 100% accuracy. This marker set could facilitate the genetic characterisation of *Macrobrachium* species in a cost-effective way for conservation and breeding purposes. However, further work is needed to validate the core SNPs identified in this study. A large sample size will have to be collected to facilitate such validation.

## Data availability

The raw genotype data for the study population is available at (10.6084/m9.figshare.17284688). The results supporting our conclusions are included in the article and the supplementary material.

## Ethical statement

No experimental animal procedures were conducted in this study.

## Author contributions

JM, EKC and FDNM conceived and developed the study design. EKC, JM, and DNT performed analyses. EKC and JM wrote the first draft of the manuscript. JM, EKC, FDNM, FS, PMO, BOO, PZ, EM, and DNT reviewed and approved the final manuscript for publication.

## Supplementary figure captions

**Supplementary Figure 1.** PCA plot obtained from a full set of 1,814 SNPs (A), 174 private SNPs (B) and 72 ‘private SNPs 80’ (C). *M_ch* – *M. chevalieri;M_dx* – *M. dux;M_fe* – *M. felicinum; M_ma* – *M. macrobrachion; M_so* – *M. sollaudii; M_sp* – *M. sp; M_vo* – *M. vollenhovenii*.

**Supplementary Figure 2.** Admixture results obtained from using 174 ‘private SNPs’ based on the ADMIXTURE software (Alexander et al., 2009).

**Supplementary Figure 3.** Admixture results obtained from using 72 ‘private SNPs’ based on the ADMIXTURE software (Alexander et al., 2009).

## References

Alexander, D.H., Novembre, J., and Lange, K. (2009). Fast model-based estimation of ancestry in unrelated individuals. Genome research 19, 1655–1664.

Bertolini, F., Galimberti, G., Calò, D., Schiavo, G., Matassino, D., and Fontanesi, L. (2015). Combined use of principal component analysis and random forests identify population-informative single nucleotide polymorphisms: application in cattle breeds. Journal of Animal Breeding and Genetics 132, 346–356.

Bowles, D.E., Aziz, K., and Knight, C.L. (2000). Macrobrachium (Decapoda: Caridea: Palaemonidae) in the contiguous United States: a review of the species and an assessment of threats to their survival. Journal of Crustacean Biology 20, 158–171.

Browder, J.A., Gleason, P.J., and Swift, D.R. (1994). Periphyton in the Everglades: spatial variation, environmental correlates, and ecological implications. Everglades: The ecosystem and its restoration, 379–418.

Cheruiyot, E., Bett, R., Amimo, J., Zhang, Y., Mrode, R., and Mujibi, F. (2018). Signatures of selection in admixed dairy cattle in Tanzania. Frontiers in genetics 9, 607.

De Grave, S., and Fransen, C. (2011). Carideorum catalogus: the recent species of the dendrobranchiate, stenopodidean, procarididean and caridean shrimps (Crustacea: Decapoda). NCB Naturalis Leiden.

Divu, D., Khushiramani, R., Malathi, S., Karunasagar, I., and Karunasagar, I. (2008). Isolation, characterization and evaluation of microsatellite DNA markers in giant freshwater prawn Macrobrachium rosenbergii, from South India. Aquaculture 284, 281–284.

Doume, C.D., Toguyeni, A., and Yao, S.S. (2013). Effets des facteurs endogènes et exogènes sur la croissance de la crevette géante d’eau douce Macrobranchium rosenbergii De Man, 1879 (Decapoda: Palaemonidae) le long du fleuve Wouri au Cameroun. International Journal of Biological and Chemical Sciences 7, 584–597.

Goudet, J. (2005). Hierfstat, a package for R to compute and test hierarchical F-statistics. Molecular Ecology Notes 5, 184–186.

Holthuis, L.B. (1980). FAO species catalogue. Volume 1-Shrimps and prawns of the world. An annotated catalogue of species of interest to fisheries.

Jin, S., Bian, C., Jiang, S., Han, K., Xiong, Y., Zhang, W., Shi, C., Qiao, H., Gao, Z., and Li, R. (2021). A chromosome-level genome assembly of the oriental river prawn, Macrobrachium nipponense. GigaScience 10, giaa160.

Jombart, T. (2008). adegenet: a R package for the multivariate analysis of genetic markers. Bioinformatics 24, 1403–1405.

Jombart, T., Devillard, S., and Balloux, F. (2010). Discriminant analysis of principal components: a new method for the analysis of genetically structured populations. BMC genetics 11, 1–15.

Kilian, A., Wenzl, P., Huttner, E., Carling, J., Xia, L., Blois, H., Caig, V., Heller-Uszynska, K., Jaccoud, D., and Hopper, C. (2012). “Diversity arrays technology: a generic genome profiling technology on open platforms,” in Data production and analysis in population genomics. Springer), 67–89.

Konan, M.K., Allassane, O., Beatrice, A.G.A., and Germain, G. (2008). Morphometric differentiation between two sympatric Macrobrachium Bates, 1868 shrimps (Crustacea: Decapoda: Palaemonidae) in West-African rivers. Journal of Natural History 42, 2095–2115.

Makombu, J.G., Oben, B.O., Oben, P.M., Makoge, N., Nguekam, E.W., Gaudin, G.L., and Brummett, R. (2015). Biodiversity of species of the genus Macrobrachium (Decapoda, Palaemonidae) in Lokoundje, Kienke and Lobe Rivers, South Region, Cameroon. Journal of Biodiversity and Environmental Science 7, 68–80.

Makombu, J.G., Stomeo, F., Oben, P.M., Tilly, E., Stephen, O.O., Oben, B.O., Cheruiyot, E.K., Tarekegn, G.M., Zango, P., and Egbe, A.E. (2019). Morphological and molecular characterization of freshwater prawn of genus Macrobrachium in the coastal area of Cameroon. Ecology and evolution 9, 14217–14233.

March, J.G., Pringle, C.M., Townsend, M.J., and Wilson, A.I. (2002). Effects of freshwater shrimp assemblages on benthic communities along an altitudinal gradient of a tropical island stream. Freshwater Biology 47, 377–390.

Matukumalli, L.K., Lawley, C.T., Schnabel, R.D., Taylor, J.F., Allan, M.F., Heaton, M.P., O’connell, J., Moore, S.S., Smith, T.P., and Sonstegard, T.S. (2009). Development and characterization of a high density SNP genotyping assay for cattle. PloS one 4, e5350.

Monod, T. (1980). In J. R. Durand, & C. Leveque (Eds.), Flore et faune aquatiques de l’Afrique sahélo-soudanienne (pp. 369–389). Paris, France: Tome I, ORSTOM.

Murphy, N.P., and Austin, C.M. (2003). Molecular taxonomy and phylogenetics of some species of Australian palaemonid shrimps. Journal of Crustacean Biology 23, 169–177.

New, M.B., Valenti, W.C., Tidwell, J.H., D’abramo, L.R., and Kutty, M.N. (2009). Freshwater prawns: biology and farming. John Wiley & Sons.

Nguyen, N.N., Kim, M., Jung, J.-K., Shim, E.-J., Chung, S.-M., Park, Y., Lee, G.P., and Sim, S.-C. (2020). Genome-wide SNP discovery and core marker sets for assessment of genetic variations in cultivated pumpkin (Cucurbita spp.). Horticulture Research 7, 1–10.

Okogwu, O.I., Ajuogu, J.C., and Nwani, C.D. (2010). Artisanal fishery of the exploited population of Macrobrachium vollenhovenii Herklot 1857 (Crustacea; Palaemonidae) in the Asu River, southeast Nigeria. Acta Zoologica Lituanica 20, 98–106.

Oliveira, R., Randi, E., Mattucci, F., Kurushima, J., Lyons, L.A., and Alves, P. (2015). Toward a genome-wide approach for detecting hybrids: informative SNPs to detect introgression between domestic cats and European wildcats (Felis silvestris). Heredity 115, 195–205.

Pakstis, A.J., Speed, W.C., Kidd, J.R., and Kidd, K.K. (2007). Candidate SNPs for a universal individual identification panel. Human genetics 121, 305–317.

Pardy, C., Motyer, A., and Wilson, S. (Year). “Resampling procedures to identify important SNPs using a consensus approach”, in: BMC proceedings: Springer), 1–6.

Pfaffelhuber, P., Grundner-Culemann, F., Lipphardt, V., and Baumdicker, F. (2020). How to choose sets of ancestry informative markers: A supervised feature selection approach. Forensic Science International: Genetics 46, 102259.

Phillips, C., Salas, A., Sanchez, J., Fondevila, M., Gomez-Tato, A., Alvarez-Dios, J., Calaza, M., De Cal, M.C., Ballard, D., and Lareu, M. (2007). Inferring ancestral origin using a single multiplex assay of ancestry-informative marker SNPs. Forensic Science International: Genetics 1, 273–280.

Powell, C. (1980). The genus Macrobrachium in West Africa. I: M. thysi, a new large-egged species from the Ivory Coast (Crustacea Decapoda Palaemonidae).

Price, A.L., Patterson, N.J., Plenge, R.M., Weinblatt, M.E., Shadick, N.A., and Reich, D. (2006). Principal components analysis corrects for stratification in genome-wide association studies. Nature genetics 38, 904–909.

Purcell, S., Neale, B., Todd-Brown, K., Thomas, L., Ferreira, M.A., Bender, D., Maller, J., Sklar, P., De Bakker, P.I., and Daly, M.J. (2007). PLINK: a tool set for whole-genome association and population-based linkage analyses. The American Journal of Human Genetics 81, 559–575.

R Core Team (2018). R: A language and environment for statistical computing. R Foundation for Statistical Computing, Vienna, Austria. URL https://www.R-project.org/.

Rosenberg, N.A., Li, L.M., Ward, R., and Pritchard, J.K. (2003). Informativeness of genetic markers for inference of ancestry. The American Journal of Human Genetics 73, 1402–1422.

Savaya-Alkalay, A., Ndao, P.D., Jouanard, N., Diane, N., Aflalo, E.D., Barki, A., and Sagi, A. (2018). Exploitation of reproductive barriers between Macrobrachium species for responsible aquaculture and biocontrol of schistosomiasis in West Africa. Aquaculture Environment Interactions 10, 487–499.

Schiavo, G., Bertolini, F., Galimberti, G., Bovo, S., Dall’olio, S., Costa, L.N., Gallo, M., and Fontanesi, L. (2020). A machine learning approach for the identification of population-informative markers from high-throughput genotyping data: application to several pig breeds. Animal 14, 223–232.

Shriver, M.D., Smith, M.W., Jin, L., Marcini, A., Akey, J.M., Deka, R., and Ferrell, R.E. (1997). Ethnic-affiliation estimation by use of population-specific DNA markers. American journal of human genetics 60, 957.

Siméon, T., Gideon, A., Dramane, D., Idrissa, C., Mexmin, K., and Pierre, N. (2014). Impact of anthropogenic activities on water quality and freshwater shrimps diversity and distribution in five rivers in Douala, Cameroon. J Bio & Env Sci 4, 183–194.

Vergamini, F.G., Pileggi, L.G., and Mantelatto, F.L. (2011). Genetic variability of the Amazon river prawn Macrobrachium amazonicum (Decapoda, Caridea, Palaemonidae). Contributions to Zoology 80, 67–83.

Weir, B.S., and Cockerham, C.C. (1984). Estimating F-statistics for the analysis of population structure. evolution, 1358–1370.

Wilkinson, S., Wiener, P., Archibald, A.L., Law, A., Schnabel, R.D., Mckay, S.D., Taylor, J.F., and Ogden, R. (2011). Evaluation of approaches for identifying population informative markers from high density SNP chips. BMC genetics 12, 1–14.

Wowor, D., Muthu, V., Meier, R., Balke, M., Cai, Y., and Ng, P.K. (2009). Evolution of life history traits in Asian freshwater prawns of the genus Macrobrachium (Crustacea: Decapoda: Palaemonidae) based on multilocus molecular phylogenetic analysis. Molecular phylogenetics and evolution 52, 340–350.

Zhang, Q.-Y., Cheng, Q.-Q., and Guan, W.-B. (2009). Mitochondrial COI gene sequence variation and taxonomic status of three Macrobrachium species. Zool Res 30, 613–619.

